# Noninvasive Cardiac Arrest Recognition Using Wearable Sensors in a Porcine Model of Ventricular Fibrillation

**DOI:** 10.64898/2026.11.18.767743

**Authors:** Zachary Englhardt, Moaz Bin Saeed, Armin Razi, Iyad Ayoub, Alvin Baetiong, Shwetak Patel, Raúl J. Gazmuri, Jake Sunshine, Vikram Iyer

## Abstract

**Background:** Wearable detection of out-of-hospital cardiac arrest (OHCA) could improve survival, given the prognostic importance of time to resuscitation and that 50% to 75% of events are unwitnessed. Existing systems may be limited by their form factor and by specificity achieved at the cost of sensitivity. We investigated whether the raw photoplethysmography (PPG) signal from low-cost wearable sensors contains a distinct signature of circulatory arrest.

**Methods:** Ventricular fibrillation (VF) was electrically induced in 9 invasively monitored pigs while raw PPG was recorded with a wearable-class sensor. PPG dynamics after VF onset were related to aortic and right atrial pressures. A heuristic detection algorithm was evaluated for sensitivity and latency by leave-one-out cross-validation in the porcine data and for false positives in 57,857 hours of free-living PPG from 115 human participants.

**Results:** After VF onset, the raw PPG showed a reproducible exponential decay (time constant 10.53 s; coefficient of variation 8.9% to 17.0% over the first 10 s) that tracked rising right atrial pressure (r = −0.90), reflecting arterial-to-venous blood redistribution. The algorithm detected VF in 8 of 9 animals (sensitivity 0.889; 95% CI, 0.518–0.997) with a median latency of 57.2 s. In the human cohort, it produced 2.73 false positives per user-year (95% CI, 1.62–4.31), comparable to current systems.

**Conclusions:** Raw wearable PPG encodes circulatory collapse during VF, as supported by invasive hemodynamic measurements. While preliminary, these results suggest the potential for wearable-based OHCA detection with improved sensitivity, reduced latency, and comparable specificity.

**Clinical Perspective:** *What Is New?:*

- After the onset of ventricular fibrillation in a porcine model, the non-pulsatile component of the wearable photoplethysmography signal, which conventional processing pipelines filter out, showed a reproducible exponential decay that tracked rising right atrial pressure.
- Adding this decay to established pulsatility-based features detected 8 of 9 arrests with a median latency of 57.2 seconds and produced 2.73 false positives per user-year across 57,857 hours of free-living human data.

*What Are the Clinical Implications?:*

- Because the sensors in current wearable devices already capture this signal, it could be made available through software changes alone and combined with existing loss-of-pulse detection to improve recognition of unwitnessed out-of-hospital cardiac arrest.

## Introduction

Out-of-hospital cardiac arrest (OHCA) remains a formidable public health challenge ^1^ because successful resuscitation depends on a sequence of rapidly executed, interdependent actions termed the Chain of Survival ^2^. The first and rate limiting link in the Chain of Survival is cardiac arrest recognition ^3^. In approximately half of OHCAs, no bystander is present and the event is unwitnessed ^4^. Thus, immediate recognition fails and every subsequent step in the Chain (calling for help, cardiopulmonary resuscitation (CPR), defibrillation, etc.) is delayed, or does not happen at all (e.g., the victim is found deceased and no resuscitation is attempted). Decades of evidence show that rapid resuscitation in the minutes following arrest improves outcomes ^5,6^.

Wearable devices present a ubiquitous potential means of OHCA recognition. While electrocardiogram (ECG) remains the gold standard for identifying cardiac arrhythmias ^7^, photoplethysmography (PPG) sensors ^8^, found in virtually all modern wearable devices, are capable of identifying cardiac arrhythmias, including those resulting in circulatory failure associated with OHCA ^9^. Shah et al. ^10^ recently demonstrated substantial progress in automated loss of pulse detection on consumer smartwatches, using machine learning algorithms to identify peripheral pulselessness with promising sensitivity and specificity, though with sensitivity that is approximately 65%. Their work confirms that wearable PPG-based detection systems could serve as a vital early identification method for sudden cardiac events. Such systems can automatically notify emergency services or nearby individuals and thus activate the Chain of Survival sooner, particularly in unwitnessed circumstances.

A central design challenge of translating these wearable systems into scalable deployments is limiting false positives; unnecessary emergency calls can overload 911 response infrastructure ^11^ and threaten the viability of wearable-based OHCA detection systems as common scenarios such as motion, sensor obstruction, and arterial occlusion (e.g., excessive pressure on the arm) can mimic loss of pulse (Fig. 1a, b). Adequate specificity has been demonstrated ^10^, but at a cost of reduced sensitivity and increased latency to classification. There is thus an opportunity to develop systems with higher sensitivity and preserved specificity without sacrificing timeliness, to pull forward the links in the Chain of Survival and improve outcomes ^5,6^ .

**Fig. 1.**
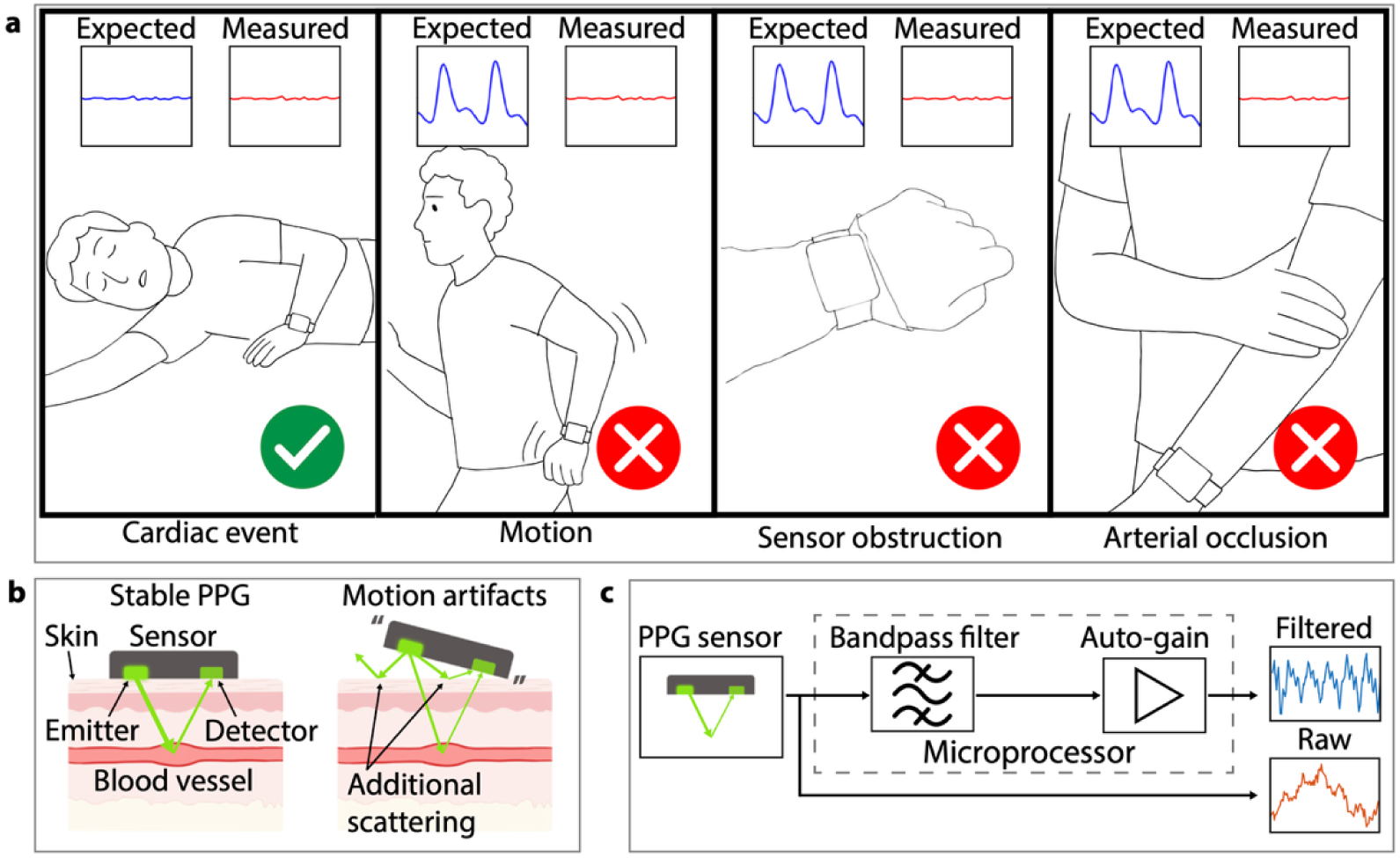
Sources of non-pulsatile PPG signals from wearable sensors. **a** Common scenarios that produce a loss of measurable pulsatility in PPG: a true cardiac event (left) versus benign causes including motion artifact, sensor obstruction, and arterial occlusion. **b** Schematic of a reflectance-mode PPG sensor under stable skin contact (left) and during motion-induced displacement (right), demonstrating how motion can result in artifacted sensor readings. **c** Standard PPG signal processing pipeline. Bandpass filtering and automatic gain control isolate the pulsatile (AC) component but remove low-frequency (DC) changes in background light absorption that may contain diagnostic information about tissue perfusion.

PPG sensors in wearable devices shine light from light-emitting diodes (LEDs) into peripheral tissue, and the reflected light is measured by a photodetector (Fig. 1b). Traditional PPG analysis has focused on the alternating current (AC) component of the signal, corresponding to pulsatile blood flow ^8^. The direct current (DC) component, or the slow-varying sub-pulsatile changes in intensity (Fig. 1c), has received considerably less attention ^8^. It is typically removed by filtering, despite potentially containing information about tissue perfusion and blood volume in the pulseless setting of OHCA.

Here, we use a porcine model ^12^ in which ventricular fibrillation (VF) is induced and allowed to persist in order to closely mimic true putative OHCA ^13–15^, a protocol that is ethically infeasible in humans. By analyzing raw green-light (537 nm) PPG from low-cost, consumer-grade sensors, we characterize PPG dynamics during the transition from normal cardiac function to persistent VF and relate them to direct measurements of aortic and right atrial pressures. We then develop a heuristics-based classifier and evaluate it on the porcine data and on a held-out human free-living dataset (115 participants, 57,857 hours), since any candidate classifier must have adequate specificity in real-world contexts.

Our findings reveal that unfiltered PPG signals exhibit distinct and reproducible exponential decay patterns during ventricular fibrillation that strongly correlate with hemodynamic collapse and blood redistribution from arterial to venous circulation, yet which are not commonly observed in real-world non-OHCA settings. These results suggest a dual approach to cardiac arrest recognition: loss of pulsatility in the AC component identifies cessation of effective cardiac output, while the novel exponential decay in the DC component reflects progressive blood redistribution from arterial to venous circulation, a hemodynamic signature not expected from other causes of non-pulsatile PPG such as motion artifact or sensor displacement.

Conditioning on both components enables robust detection on porcine model data as well as improved false-positive rejection on human free-living data, suggesting the utility of wearable PPG-based monitoring algorithms that leverage the full, unfiltered signal.

## Methods

### Study design and animal model

The animal study was approved by the Institutional Animal Care and Use Committee at Rosalind Franklin University of Medicine and Science (protocol #23-20) and was conducted according to institutional guidelines. This was an exploratory, observational study in a porcine model of electrically induced VF. Ten male domestic pigs (35–40 kg) were studied. Successful VF induction was the a priori criterion for inclusion; induction was unsuccessful in the first animal, which was excluded, leaving nine analyzed experiments (n = 9). Each animal served as its own control, with its pre-VF baseline compared against the post-VF period. No formal a priori sample-size calculation was performed.

### Instrumentation and experimental protocol

Animals were sedated with intramuscular ketamine, intubated under intravenous propofol, maintained on isoflurane anesthesia, placed in a supine position, and instrumented with multiple physiological sensors (Fig. 2a). Each was equipped with a custom wearable-class PPG sensor on the ear and hoof. Additional sensors included a limb-mounted ground-truth clinical PPG sensor, ECG for cardiac rhythm monitoring, a CO2 sensor, and surgically implanted fluid-filled catheters to continuously measure pressures in the right atrium, left ventricle, pulmonary artery, and aorta.

**Fig. 2.**
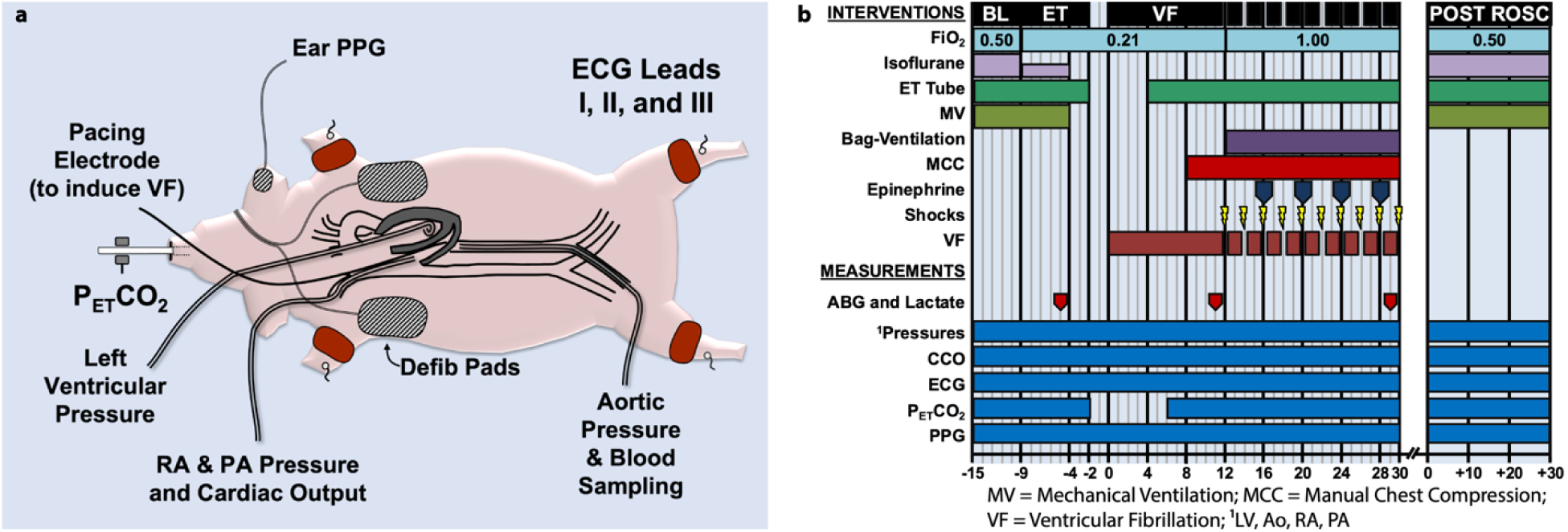
Porcine model of ventricular fibrillation with simultaneous PPG and hemodynamic monitoring. **a** Sensor placement and invasive instrumentation. Animals were equipped with a wearable PPG sensor on the ear, ECG leads, a CO2 sensor, a right ventricular pacing electrode, defibrillation pads, and fluid-filled catheters for continuous measurement of aortic, right atrial, left ventricular, and pulmonary artery pressures. PPG, photoplethysmography; PETCO2, end-tidal PCO2; RA, right atrium; PA, pulmonary artery. **b** Experimental protocol timeline showing the sequence of interventions (fraction of inspired oxygen, isoflurane anesthesia, endotracheal intubation, mechanical and bag ventilation, manual chest compression, epinephrine, defibrillation shocks, and ventricular fibrillation) alongside the continuous measurements (arterial blood gas and lactate sampling, intravascular pressures, continuous cardiac output, ECG, end-tidal CO2, and PPG) across the baseline (BL), extubation (ET), untreated ventricular fibrillation (VF), and post–return-of-spontaneous-circulation (POST ROSC) phases. MV, mechanical ventilation; MCC, manual chest compression; CCO, continuous cardiac output; ROSC, return of spontaneous circulation.

Fig. 2b illustrates the experimental protocol. Animals were extubated following confirmation of stable spontaneous breathing under anesthesia. VF was then electrically induced via a pacing electrode in the right ventricle and left untreated for 10 minutes in the first three analyzed animals and 8 minutes in the subsequent six, after which standard resuscitation protocols were initiated. All quantitative PPG analyses are confined to the first 200 seconds after VF onset, before this difference applies. VF onset was determined from the ECG independently of the PPG signal. Data was recorded continuously from all sensors throughout the protocol, except for end-tidal CO2 (PETCO2), which ceased upon extubation. Surgical instrumentation, anesthesia management, VF induction, and resuscitation procedures are detailed in the Supplemental Methods.

### Wearable PPG sensor

Raw PPG was acquired with a MAX30101 sensor (Analog Devices) using the green LED channel (537 nm) at a nominal sampling rate of 36 Hz. The raw green-channel intensity was used for all analyses. Sensor hardware, firmware, and data logging are described in the Supplemental Methods.

### Human free-living dataset

False positives were evaluated on a previously described free-living dataset of raw PPG from 120 participants (60,853 hours)^16^. The data were collected under a protocol approved by the Institutional Review Board at the University of Memphis, which included informed consent as described in the accompanying publication. Data was obtained from its authors according to their data access protocols. All recordings from five participants (2,996 hours) were reserved for parameter tuning and excluded from evaluation, leaving 115 participants (57,857 hours) in the held-out evaluation cohort. The dataset contains no known VF events.

### Detection pipeline

PPG waveforms were resampled to 25 Hz and evaluated by a four-stage pipeline of six gain-invariant algorithms. Five algorithms compute frequency and signal entropy features, and the sixth, the decreasing threshold, identifies sustained monotonic decreases in the DC component. A candidate detection is emitted when all algorithms meet their trigger counts within a sliding 4-second window, and is confirmed only if comparisons between a pre-decline baseline window and the window preceding detection are satisfied. Algorithm thresholds were selected by grid search within leave-one-out cross-validation across the nine porcine experiments, using the training animals together with the five-participant human tuning set. For evaluation on the human cohort, the selection was repeated using all nine porcine experiments. Algorithm definitions, segment-comparison constraints, and the parameter selection procedure are detailed in the Supplemental Methods.

### Statistical analysis

All analyses were performed using custom MATLAB (R2024b, MathWorks) and Python scripts. Group data are reported as mean ± standard deviation unless otherwise noted, and n denotes the number of porcine experiments (n = 9). For the pressure–PPG correlation analysis, Pearson correlation coefficients were computed for each experiment between the normalized raw ear PPG signal and the simultaneously recorded right atrial or aortic pressure within each time window relative to VF onset. Per-experiment coefficients were aggregated across the nine experiments using Fisher z-transformation to obtain the mean correlation and its 95% confidence interval based on the between-experiment variance, an approach that stabilizes the variance of correlation estimates and is better suited to the small sample size than averaging raw coefficients. The statistical significance of each correlation was assessed with a two-tailed one-sample t-test evaluating whether the mean of the per-experiment Fisher z-transformed coefficients differed from zero (n = 9 experiments, df = 8), with α = 0.05; exact P values are reported in the Results.

For the detection pipeline, sensitivity on the porcine data was estimated by leave-one-out cross-validation across the nine experiments, with the 95% confidence interval computed using the Clopper–Pearson exact binomial method, and detection latency is reported as the median and interquartile range over the 8 detected events. The false-positive rate on the human free-living cohort (115 participants, 57,857 hours) is reported per user-year with a 95% confidence interval computed using the exact Poisson method. Algorithm thresholds and the ramp-start tolerance were selected by grid search within each cross-validation fold (comparison constants were fixed a priori), and the final human-cohort evaluation used parameters fit on all nine porcine experiments.

## Data availability

All data necessary for interpreting the manuscript has been included. Porcine wearable PPG data is available on Dryad (<u>temporary reviewer sharing link</u>).

We are unable to redistribute the third-party human PPG dataset^16^ ourselves. To inquire about access to this third-party data, please contact the MOODS team at.

## Code availability

Code including the detection pipeline, the leave-one-out cross-validation, and ablations, is available on Zenodo (temporary reviewer sharing link).

## Results

### Confirmation of ventricular fibrillation and expected PPG response

Fig. 3a shows the raw PPG output beginning from pre-VF baseline monitoring through approximately 7.5 min after VF induction. Fig. 3b-d shows the PPG and ECG signals with higher resolution in short time windows before and after VF. During the baseline period, we observe a normal sinus rhythm in the ECG, and corresponding periodic PPG waveforms from both the wearable and ground truth PPG. After VF induction, we confirm disorganized, rapid ventricular electrical activity indicative of VF via ECG (Fig. 3d).

**Fig. 3.**
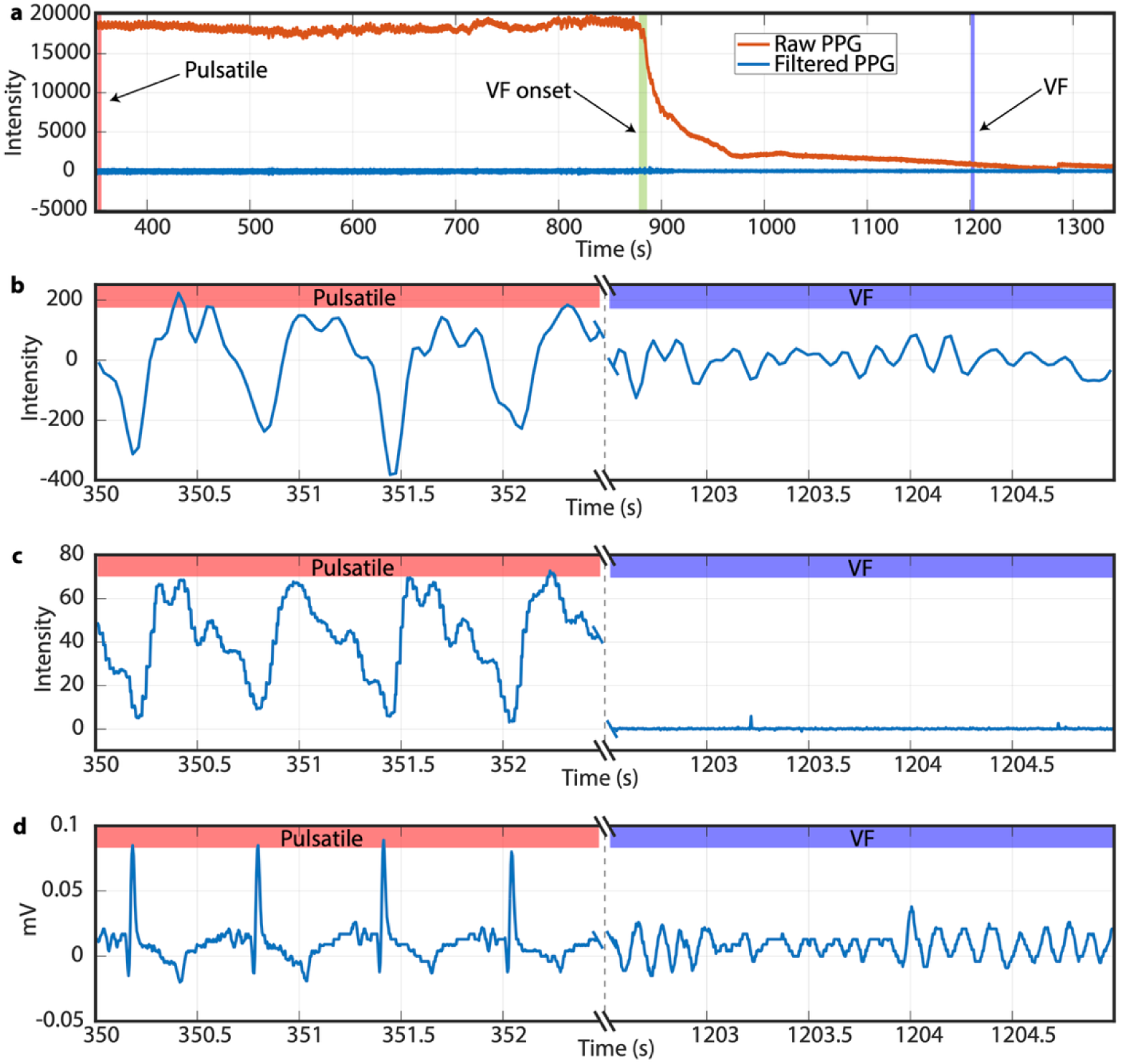
Raw PPG reveals DC-component changes at VF onset not visible in filtered signals. **a** Full-duration ear PPG recording spanning baseline through post-VF onset. The raw (unfiltered) signal (orange) exhibits a marked decrease in DC offset following VF induction; the bandpass-filtered signal (blue, 0.25–10 Hz) shows only the expected loss of pulsatility since DC offset is removed. Shaded markers indicate the segments expanded in b–d and the point of VF induction. **b** Representative 2.5-second segments of bandpass-filtered wearable PPG during baseline (left) and VF (right), showing loss of the periodic pulsatile waveform. **c** Corresponding segments from a clinical PPG sensor with built-in hardware filter, confirming comparable waveform morphology to filtered wearable sensor. **d** Corresponding ECG segments showing normal sinus rhythm before VF induction and disorganized ventricular activity during VF.

During baseline phases, we compare the PPG from the wearable sensor and the ground-truth medical data acquisition system. Since the wearable sensor signal is unfiltered, we applied a sixth order Chebyshev type II filter with a stopband attenuation of 40 dB and a passband between 0.25 Hz and 10 Hz to extract the pulsatile waveform to enable comparison with the pre-filtered signal from the clinical PPG sensor. The filter effectively removes the slow varying DC signal changes (below 0.25 Hz) and smooths the signal by removing high frequency noise above 10 Hz. Filter parameters are chosen based on the findings of Liang et al. ^17^ which describes optimal PPG filter parameters.

During baseline phases, both the ground truth PPG and the bandpass-filtered mobile sensor PPG signals exhibit the expected pulsatile waveform morphology with regular and consistent pulse patterns prior to VF induction. Following VF induction, both signals lose discernible pulsatility, consistent with prior reports on PPG during VF ^18–21^: the clinical trace becomes near-flat, while the filtered wearable trace retains only low-amplitude, aperiodic residual activity (Fig. 3b, c).

### Analysis of raw PPG waveform

Analysis of the raw PPG signal shown in Fig. 3a reveals a distinct alteration in its waveform at VF onset while the filtered PPG remains flat. This is observed as a marked decrease in the DC component of the waveform, decaying rapidly in intensity with the loss of pulse. A new steady state is approached after approximately one minute. Crucially, this behavior is not observable in the signal when typical PPG data processing steps such as band-pass filtering and automatic gain control are applied, as the decrease occurs at a much longer timescale than individual heartbeats. Thus, our subsequent analysis will leverage the mobile PPG sensor where we can directly access raw (unfiltered) sensor readings that contain the complete spectrum of both AC and DC PPG components.

### Aggregate PPG response across subjects

Before examining the correlation between the observed PPG signals and circulatory system behavior, we evaluate the consistency of this PPG response across all 9 experiments. After time-alignment to the VF onset point and normalization of signal amplitudes, we calculate the mean and standard deviation of each raw PPG response as shown in Fig. 4a. To account for the difference in the baseline amplitude measured by the PPG signal between subjects, we scale the mean measured value in the 20 seconds prior to VF to a value of 100%, with a value of 0% corresponding to a value 0 on the sensor.

**Fig. 4.**
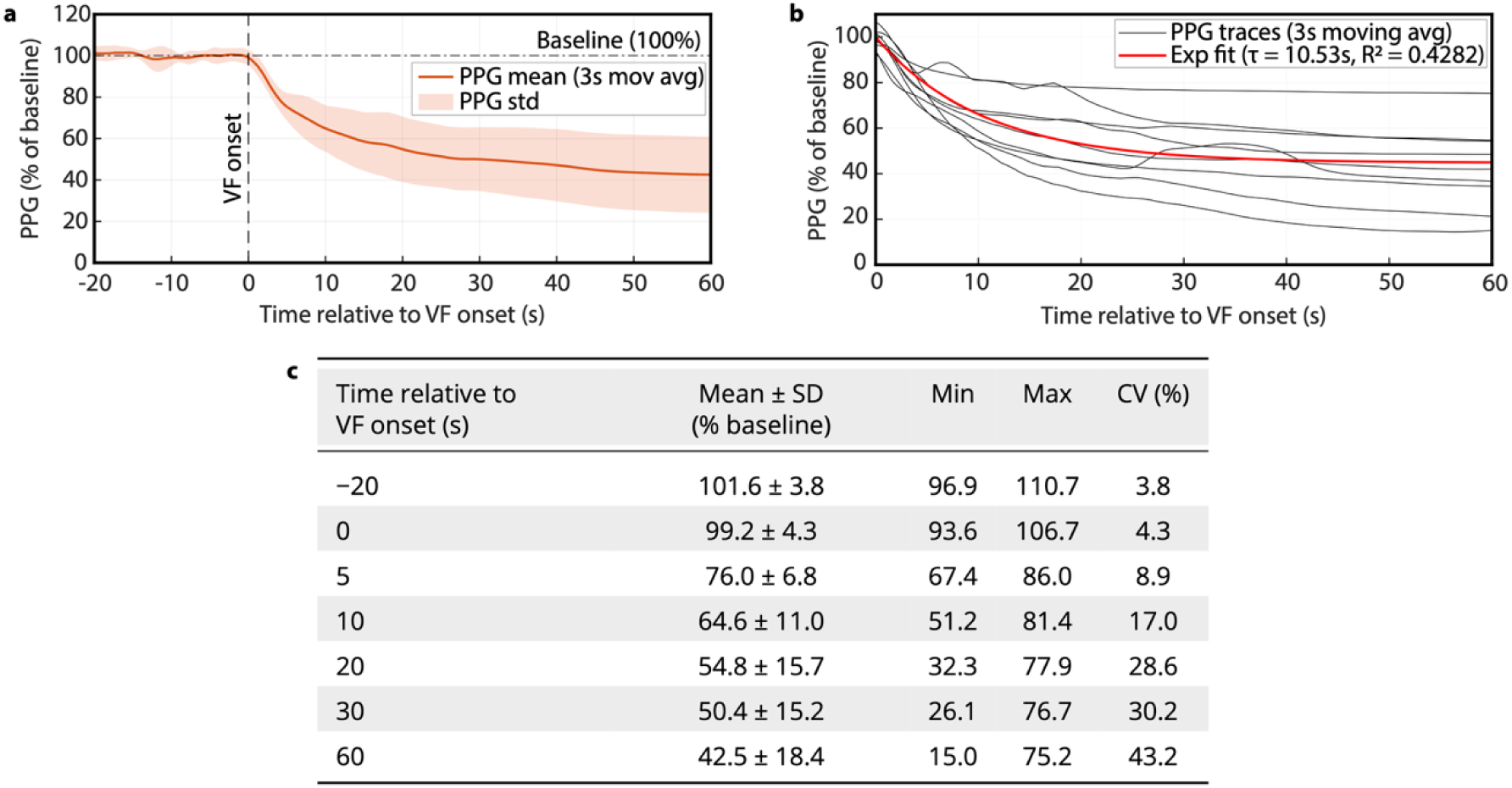
Exponential decay of the DC component of PPG following VF onset is consistently observed across subjects. **a** Mean and standard deviation of normalized ear PPG across 9 experiments as a function of time relative to VF onset (3-second moving average). Solid line, mean; shaded region, standard deviation. Signal amplitudes are normalized to the mean of the 20 seconds preceding VF (100%). **b** Single-exponential model, y(t) = 100 + b(1 − e^−t/τ^), fitted to the nine individual ear PPG traces pooled over 0–60 s after VF onset (τ = 10.53 s, R² = 0.4282), overlaid on those traces (3-s moving average, normalized as in a). **c** Mean ± SD, minimum, maximum, and coefficient of variation of wearable PPG intensity, expressed as a percentage of the pre-VF baseline, at selected time points; inter-subject variability is low through the first 10 s (CV 8.9% at 5 s, 17.0% at 10 s) and the maximum across subjects falls monotonically, so every animal is below baseline at every time point after VF onset.

To characterize the temporal dynamics of this PPG response, we developed an exponential decay model to capture the signal degradation following cardiac arrest. As shown in Fig. 4b, we fit a single-exponential model, y(t) = 100 + b(1 − e^−t/τ^), to the normalized DC components of all 9 subjects pooled over the 60 s following VF onset, yielding a time constant τ = 10.53 s (R² = 0.4282, evaluated against the pooled individual traces). The agreement between the model and the experimental data indicates that this simple exponential function captures the shared early decay across animals, with the moderate R² reflecting the between-animal spread in steady-state level visible in Fig. 4b.

Critically, Fig. 4b shows that each signal exhibits rapid decay in the first five seconds, and Fig. 4c quantifies the consistency of this decay across subjects over the first 10 seconds, with a low coefficient of variation (8.9% at 5 s, 17.0% at 10 s), followed by settling to different relative steady-state values. Despite inter-subject physiological differences, the signal dynamics remain consistent during the critical early phase of circulatory collapse, and over the 60 s following VF onset, every subject experiences a decrease in normalized PPG intensity relative to pre-VF levels. This combination of low early-phase variability and an approximately exponential decay in combination with loss of pulsatility could act as a novel feature supporting differentiation between true cardiac arrest and confounding conditions (such as those shown in Fig. 1a).

### Relationship to aortic and right atrial pressures

Because the aorta represents the beginning of the arterial system and the venous system terminates in the right atrium, we recorded aortic pressure (AOP) and right atrial pressure (RAP) concurrently with the raw ear PPG. In an individual experiment (Fig. 5), both pressure waveforms are pulsatile and follow the sinus rhythm of the ECG before VF, and change at the same time as the raw PPG following VF induction (Fig. 5d). AOP loses its pulsatile profile and falls from a pre-VF mean of 109.1 mmHg to 21.1 mmHg at 20 s. RAP also loses pulsatility but rises from 7.5 mmHg to 14.1 mmHg at 20 s, with a transient peak of 25.0 mmHg. By 60 s the two pressures have converged (AOP 20.0 mmHg, RAP 19.7 mmHg).

**Fig. 5.**
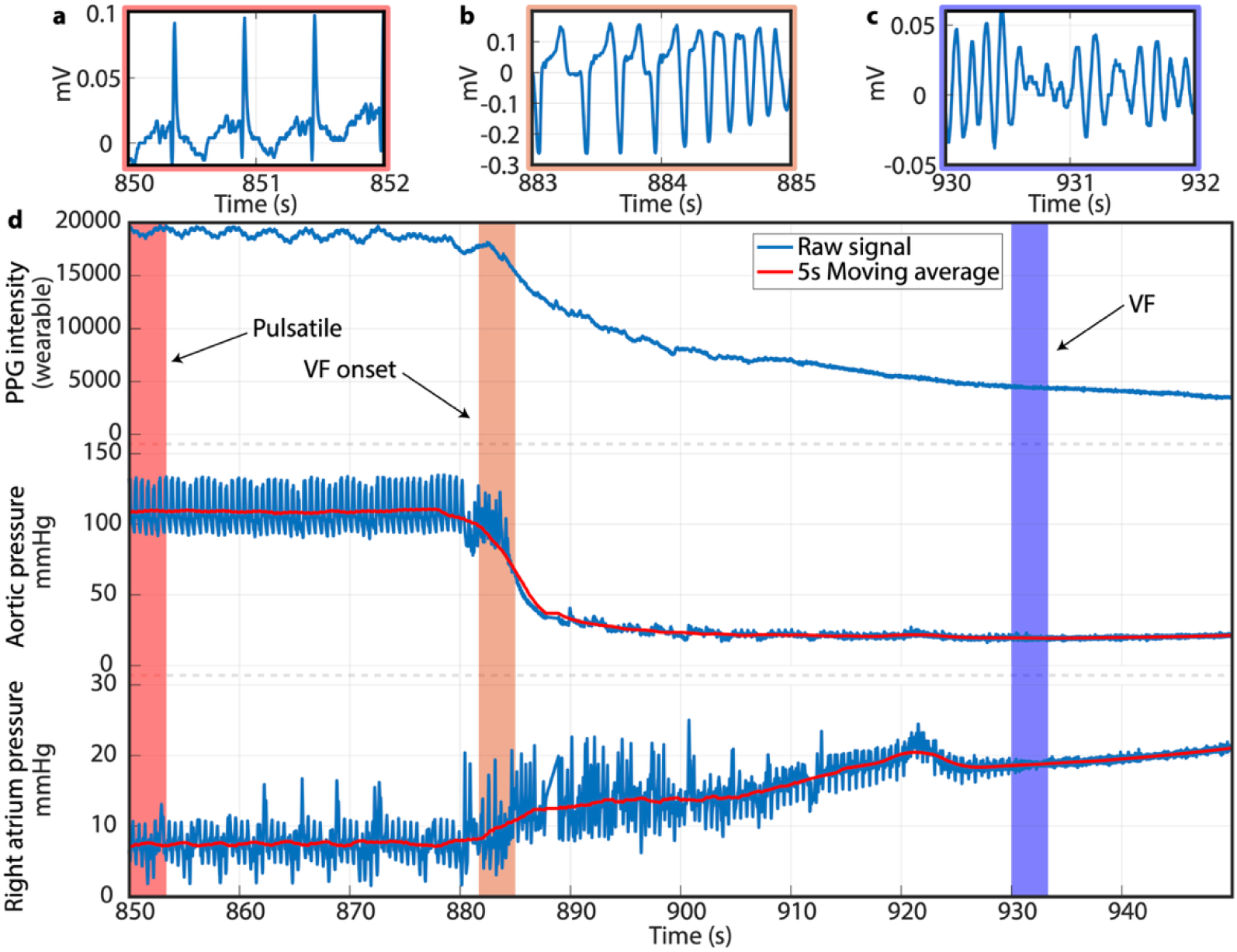
Temporal correspondence between raw PPG and invasive hemodynamic measurements during VF. **a–c** Representative ECG snapshots confirming cardiac rhythm state: normal sinus rhythm before VF (**a**), transition to VF (**b**), and established VF (c). **d** Simultaneous recording of unfiltered ear PPG, aortic pressure, and right atrial pressure. The DC component of PPG declines in parallel with falling aortic pressure and rising right atrial pressure following VF onset. Blue traces, raw signal; red traces, 5-s moving average. Shaded bands mark the intervals shown in a (red), b (orange), and c (purple).

Across all nine experiments, PPG appears negatively correlated with RAP and positively correlated with AOP (Fig. 6a, b). Pearson correlation coefficients were calculated between each raw PPG signal and the corresponding pressure waveforms, with confidence intervals for the mean coefficients determined using Fisher’s Z-transformation^22^. A strong negative correlation was observed between PPG and RAP over the full 60 s after VF onset (r = -0.90, p = 5.8 × 10⁻⁷) and within each window [0–10 s (r = -0.91, p = 1.6 × 10⁻⁵), 10–30 s (r = -0.85, p = 2.3 × 10⁻⁶), and 30–60 s (r = -0.91, p = 0.0035)] (Fig. 6c). PPG and AOP were strongly correlated over the full window (r = 0.85, p = 1.3 × 10⁻⁸), but only in the initial 0–10 s window (r = 0.96, p = 8.2 × 10⁻⁶), with no significant correlation at 10–30 s (r = 0.59, p = 0.089) or 30–60 s (r = -0.38, p = 0.42).

**Fig. 6.**
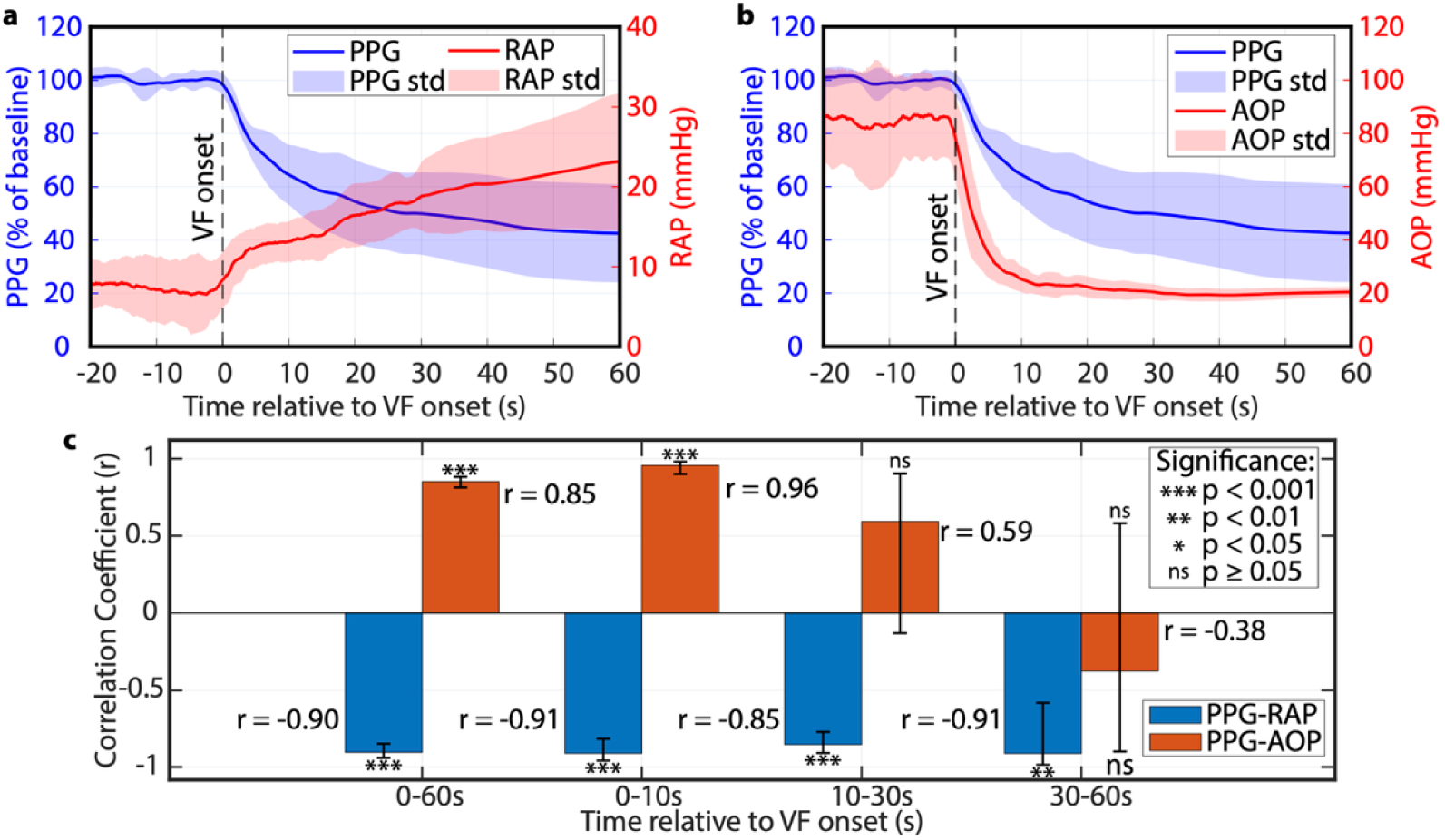
The DC component of PPG correlates with venous pressure throughout circulatory collapse. **a, b** Mean and standard deviation of normalized ear PPG (n = 9) plotted against right atrial pressure (**a**) and aortic pressure (**b**) as a function of time relative to VF onset (3-second moving average). Solid lines, means; shaded regions, standard deviations. **c** Pearson correlation coefficients between raw ear PPG and right atrial and aortic pressures across successive time windows following VF onset. Bars show the mean correlation across the nine experiments computed via Fisher z-transformation; error bars denote the corresponding 95% confidence intervals. The PPG signal maintains a strong negative correlation with right atrial pressure across all intervals (r = -0.85 to -0.91, all p < 0.01), whereas the positive correlation with aortic pressure is significant only during the first 10 seconds. In c, three asterisks indicate p < 0.001, two asterisks p < 0.01, one asterisk p < 0.05, and ns denotes not significant (p ≥ 0.05); exact P values are reported in the main text.

We hypothesize that this RAP-dominant behavior reflects the hemodynamic transformation during VF. Cessation of coordinated contraction eliminates the pressure gradient between the arterial and venous circulations ^23^, and blood redistributes passively from the high-pressure arterial system to the low-pressure venous compartment, slowed by the resistance of the circulatory network ^24^, until pressures nearly equilibrate. Across the nine experiments at 55–60 s after VF onset (mean ± SD), RAP reached 22.86 ± 8.24 mmHg while AOP decreased to 20.34 ± 1.98 mmHg (the RAP spread reflects one animal at 44.3 mmHg; the other eight ranged from 18.3 to 24.1 mmHg). That PPG remains correlated with RAP throughout the post-VF period while losing correlation with AOP suggests the DC component primarily reflects venous engorgement of the tissue, consistent with the proximity of venous vessels to the skin surface and their direct hydraulic connection to the right atrium.

### Developing a prototype classifier

Our motivation is to detect cardiac arrest in free-living scenarios, not controlled hospital settings. End-to-end training and evaluation on human free-living data would require labels of true VF events, which is unfeasible. Instead, we developed an algorithm using our porcine data and recordings from a held-out set of five human tuning participants, then evaluated false positives on 57,857 hours of raw PPG from the remaining 115 participants of a 120-participant free-living cohort ^16^.

Two factors shaped the design. First, mean PPG intensity differs by approximately two orders of magnitude between the porcine and human datasets, reflecting differences in sensor gain, optical coupling, and hardware, although the 95% confidence intervals of pulsatile-band (0.25–10 Hz) signal-to-noise ratio overlap (Fig. 7a). This precludes approaches that rely on absolute amplitude, including supervised classifiers on raw features, which could trivially separate the populations by scale, and requires every feature to be self-normalized and amplitude-invariant. Second, the human data contain no known VF events, so a trained model could not be distinguished from the naive baseline of never detecting VF.

**Fig. 7.**
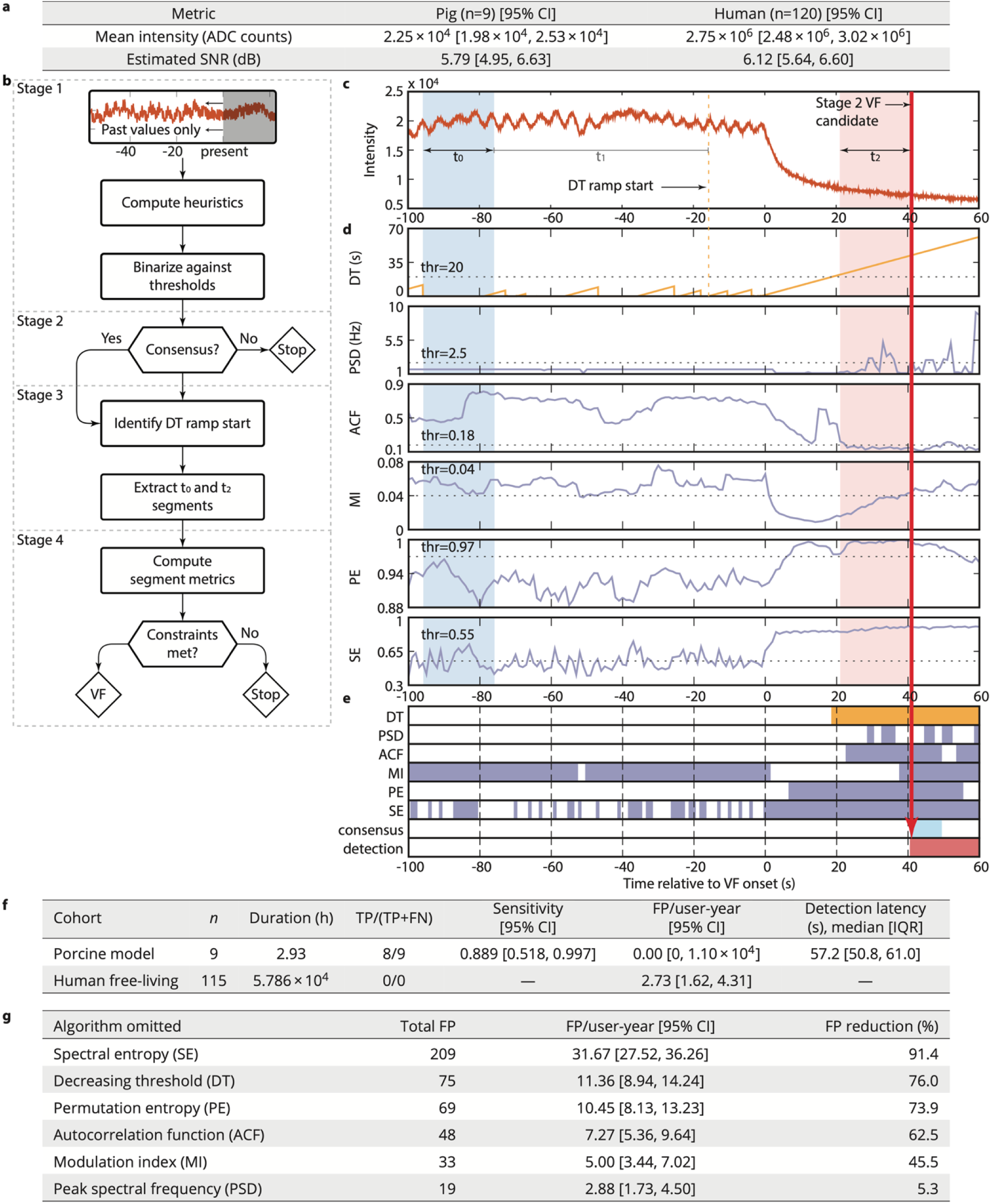
Gain-invariant detection pipeline and evaluation on porcine and human free-living data. **a** Comparison of mean PPG intensity and pulsatile-band signal-to-noise ratio (SNR) between porcine and human datasets; absolute amplitudes differ by approximately two orders of magnitude while SNR confidence intervals overlap. **b** Schematic of the four-stage detection pipeline: per-algorithm triggering against thresholds (Stage 1), windowed consensus requiring all algorithms to fire (Stage 2), identification of the decreasing threshold ramp onset (stepping over interruptions of the decrease no longer than the ramp-start tolerance) and extraction of baseline (t₀) and recent (t₂) segments (Stage 3), and segment-comparison constraints (Stage 4). **c** Representative raw ear PPG trace from a single porcine experiment, plotted relative to VF onset, showing the decreasing-threshold ramp start located in Stage 3 (dashed line), the segment windows (t₀, t₁, t₂), and the Stage 2 VF candidate (red line) that is confirmed by the segment comparison. **d** Corresponding per-algorithm outputs with threshold values (dotted lines). Algorithms derived from prior literature (purple) operate on pulsatile signal features; the decreasing-threshold algorithm (orange) operates on the DC component. **e** Binary trigger state for each algorithm, the Stage 2 consensus candidate, and the confirmed detection. **f** Detection performance on porcine data (n = 9) and human free-living data (n = 115, 57,857 hours). 95% confidence intervals computed via Clopper-Pearson (sensitivity) and exact Poisson (FP rate). Detection latency is summarized as the median and IQR over the 8 detected porcine events; duration for the porcine cohort is the cumulative pre-VF monitoring time (2.93 h) over which false positives could occur. **g** Leave-one-algorithm-out ablation on human data showing the effect of individually removing each algorithm on false-positive count; percent reduction indicates each algorithm’s contribution to specificity within the full ensemble. Panel a characterizes the full free-living cohort (120 participants, 60,853 h), whereas panels f and g report evaluation on the 115-participant held-out set (57,857 h) that remained after all data (2,996 h) from the 5 participants used for parameter tuning were excluded.

We therefore designed a multi-algorithm pipeline in which each metric is a gain-invariant heuristic, allowing the same features and parameters to be applied to porcine and human PPG (Fig. 7b; Methods). Six algorithms compute metrics on trailing PPG windows aligned to a 1-second grid (Stage 1). A candidate detection is emitted when all algorithms meet their trigger counts within a sliding 4-second window (Stage 2). The onset of the preceding DC decline is then used to place a pre-decline baseline window (Stage 3), and the candidate is confirmed only if comparisons between this baseline and the window preceding detection are satisfied (Stage 4). A 300-second refractory period suppresses repeat triggers.

Five algorithms evaluate frequency and signal entropy features drawn from prior PPG signal processing features ^25–28^. The sixth, the decreasing threshold (DT), is novel and designed to capture the DC decay identified above. Because exponential fit parameters are substantially affected by noise and the choice of start time^29^, the DT test instead identifies sustained monotonic decreases in the DC component without requiring an explicit onset estimate. By operating on the DC portion of the signal, this test measures a feature mechanistically distinct from existing approaches. Fig. 7c–e shows a porcine PPG excerpt at VF onset, the corresponding heuristic values, per-algorithm trigger states, and the resulting detection.

### Classifier evaluation

We evaluated the detection algorithm on the porcine data using a leave-one-out cross-validation scheme ^30,31^. Each fold trains on 8 pigs; in each of the nine folds, the six Stage 1 thresholds and the Stage 3 ramp-start tolerance are selected by a grid search using the training pigs together with a fixed tuning set of 5 human participants that is excluded from the human evaluation cohort (see Methods). The resulting algorithm is then applied to the remaining pig PPG trace. VF is detected in 8 of 9 folds yielding a sensitivity of 0.889 (95% CI [0.518, 0.997]), median latency of 57.2 s (interquartile range [IQR] 50.8–61.0 s; n = 8), and no false positives as shown in Fig. 7f.

For evaluation on the human free-living dataset, thresholds were recomputed with no held-out pigs and tested on the 115 participants (57,857 h) that remained once all data (2,996 h) from the 5 parameter-tuning participants were excluded from the full 120-participant cohort (60,853 h); a fixed subset of those participants’ recordings (618 h after gap splitting) was scored during parameter selection. The full ensemble pipeline produced a total of 18 false positives across 15 participants, corresponding to a rate of 2.73 (95% CI [1.62, 4.31]) per user-year (Fig. 7f). We note that latency and sensitivity cannot be estimated as there are no true positive VF events in the human cohort.

Fig. 7g reports, for each leave-one-algorithm-out variant of the pipeline, the false-positive count, the rate per user-year with an exact Poisson 95% confidence interval, and the percent reduction in false positives achieved by the full pipeline relative to that variant. False-positive reductions range from 5.3% to 91.4%. For five of the six ablations, the full pipeline produces at least 45% fewer false positives than the corresponding ablated pipeline, consistent with complementary contributions at the selected operating point; omitting peak spectral frequency produces the smallest difference (5.3%).

Removing the decreasing-threshold consensus requirement while retaining DT-derived baseline placement raises the false-positive count from 18 to 75, corresponding to 76.0% fewer false positives with the full pipeline, the second-largest single contribution in the ensemble after spectral entropy, equivalent to a roughly 4-fold reduction in false-positive rate from 11.36 [8.94, 14.24] per user-year without it to 2.73 [1.62, 4.31] per user-year for the full pipeline. That the decreasing threshold test continues to provide substantial false-positive rejection in the presence of the five other signal features indicates that the DC component of PPG carries discriminative information useful in addition to the other features evaluated here. This comparison is conservative as it measures the incremental contribution of the DT consensus requirement, with the ablated condition without DT consensus still benefiting from DC-derived baseline comparison localization. Taken together, the porcine and human evaluations suggest that the DC-decay dynamics identified in this work represent a complementary signal feature for cardiac arrest detection, one that future systems could integrate alongside AC-domain features within more expressive modeling frameworks.

## Discussion

Wearable PPG sensors, now ubiquitous in consumer devices, present a scalable opportunity to detect critical emergencies such as OHCA. While traditional PPG processing pipelines isolate the pulsatile AC signal for heart rate and oxygen saturation calculations ^26–28^, we show this can remove information that improves detection specificity. In a porcine model designed to closely mimic OHCA, the unfiltered PPG exhibits a distinct and reproducible exponential decay (τ ≈ 10.5 s) following VF onset.

This decay likely reflects fundamental hemodynamic changes of cardiac arrest. With the onset of VF, the heart ceases to generate arterial pressure or remove blood from the venous system; blood redistributes from the arterial to the venous circulation and pressures equalize. The increasing blood volume in peripheral venous beds, particularly in superficial tissue, leads to greater absorption of the sensor’s emitted light. Our correlation analysis supports this explanation: PPG correlates negatively with RAP throughout the post-VF period and positively with AOP only during the first 10 s. Such redistribution is not expected during benign, everyday events that reduce PPG signal strength.

By combining existing PPG features with this insight into difficult-to-obtain post-VF physiology, our classifier achieves a porcine sensitivity (0.889, 95% CI [0.518, 0.997]) that, if reproduced in humans, would place this approach on comparable footing with existing consumer-device systems while relying on PPG alone, without inertial or user-prompt modalities. On the human free-living cohort, it achieved an acceptably low false-positive rate (2.73 per user-year, 95% CI [1.62, 4.31]), comparable to that reported by Shah et al. ^10^, where similar false detection frequencies were mitigated through user interface design. The DC-based heuristic contributes a roughly 4-fold reduction in this rate (Fig. 7g).

Several limitations should be considered. The porcine cohort is small (n = 9), yielding a wide confidence interval on sensitivity, and the protocol examined only VF; other arrest mechanisms may produce different DC-component dynamics. The controlled laboratory setting does not capture variability inherent to ambulatory humans, including posture, skin pigmentation, body composition, and sensor coupling. Sensitivity and latency are estimated only from porcine data and false-positive rate only from human data, as neither dataset permits evaluation of both.

Finally, the heuristic pipeline reflects the constraints of small, event-limited datasets rather than an assertion that heuristic methods are optimal for production deployment. Regardless of whether the specific DC decay mechanism identified here generalizes fully to human physiology, these findings nonetheless underscore the potential value in recording unfiltered PPG signals in both research and clinical data collection efforts.

That these results use low-cost sensors at the most common wavelength (green, 537 nm) points to an opportunity to make this functionality accessible on lower-cost platforms. Future research should confirm the DC dynamics and classifier sensitivity on true-positive VF events in humans, and combine the DC metric with proven modalities such as inertial motion detection and user notifications, which reduce false positives^10^. More broadly, unfiltered PPG is already available on some consumer devices, including those used in our free-living cohort, and can be exposed on others with straightforward software changes, creating opportunities to detect other cardiac conditions and to explore different sensor placements (e.g., wrist, ear, finger). Collectively, these findings seek to advance the transformation of wearables from wellness-oriented fitness trackers into clinically actionable, potentially life-saving monitoring tools.

## Acknowledgments

We thank Qiuyue (Shirley) Xue for initial sensor prototypes and Mithun Saha and Maxwell A. Xu for their assistance in onboarding the human PPG dataset. Large language model (LLM)-based tools were used to assist with writing and debugging the data-analysis and figure-generation programs and to proofread the manuscript. All AI-assisted code and text were reviewed, verified, and edited by the authors, who take full responsibility for the content and for the accuracy of the reported analyses and results.

## Supplemental Material

Supplemental Methods

